# Digital Atlases to Unlock the Potential of Brain Biorepository Tissues for Interdisciplinary Research

**DOI:** 10.64898/2026.05.13.724753

**Authors:** Jason M. Webster, Ali Shojaie, Yiqin Alicia Shen, Tung Le, Emily Ragaglia, Marika Bogdani, Amanda Kirkland, Christine Mac Donald, Caitlin S. Latimer, C. Dirk Keene, Thomas J. Grabowski

**Affiliations:** Integrated Brain Imaging Center, Department of Radiology, University of Washington, Seattle, WA, USA; Department of Biostatistics, University of Washington, Seattle, WA, USA; Department of Laboratory Medicine and Pathology, Division of Neuropathology, University of Washington, Seattle, WA, USA; Department of Neurological Surgery, University of Washington, Seattle, WA, USA

## Abstract

Human brain tissue preserved in biorepositories is foundational for the structural, cellular, and biomolecular research necessary for a mechanistic understanding of neurological diseases. Realizing the research potential of these valuable resources requires well-characterized research-relevant tissue that can be efficiently identified by investigators and incorporated into the conceptual and computational frameworks of interdisciplinary research. Several large-scale efforts to improve research reliability and reproducibility have sought to characterize and annotate the processes by which these samples are collected, yet limited progress has been made on standardizing spatial information for these samples. Biorepositories systematically collect brain tissue according to a brain sampling protocol (BSP) that differs between institutions, yet explicit spatial information regarding the samples may not be documented in standard operating procedures (SOPs). The amount of anatomical location details available to investigators are inconsistent across biorepositories and typically lack sufficient anatomical precision to ensure correspondence with samples from other biorepositories or research relevant brain regions specified by neuroimaging, functional, or disease-susceptibility criteria. Here, we introduce a pipeline for developing a Spatial Atlas for Mapping Protocol Locations of Ex vivo Samples (SAMPLES), which uses a neuroimaging framework to create a 3D representation of a BSP through a metrically precise digital instantiation of the procedures for brain extraction, segmentation, slicing, and sampling on a modern digital brain template. SAMPLES incorporates modern neuroinformatics conventions to create explicit 3D labels of BSP-defined samples that can be interactively visualized with freely available neuroimaging software. We illustrate the pipeline by developing an atlas for the protocol from the University of Washington BioRepository and Integrated Neuropathology laboratory (UW BRaIN SAMPLES). By providing an explicit, computable reference, SAMPLES atlases can support the efficient identification, referencing, and utilization of postmortem samples for interdisciplinary research. These capabilities enable biorepository workflows, data harmonization across biorepositories, and integration with antemortem neuroimaging.

## Introduction

Neurological conditions have recently overtaken cardiovascular diseases as the leading cause of global health loss (Steinmetz et al., 2024). Brain tissue is essential for elucidating the cellular and molecular mechanisms necessary for the development of effective treatments and interventions for a wide range of neurological diseases (Murray et al., 2025; Palmer-Aronsten, Sheedy, McCrossin, & Kril, 2016). A fundamental requirement of this research is the accessibility of human brain tissue systematically collected and preserved in biorepositories. The procedures by which postmortem brain tissue is prepared, sectioned, sampled, processed, and preserved constitute a brain sampling protocol (BSP). These BSPs can differ across diagnoses, over time, or between biorepositories (Danner et al., 2024; Lucot et al., 2023). Many successful and ongoing efforts have improved the reliability and reproducibility of research using brain tissue through standardizing, characterizing, and annotating the processes by which brain tissue samples are collected for specific protocols (LaBaer, Miceli, & Freedman, 2018; Moore et al., 2011; Ravid & Park, 2014). However, an explicit neuroinformatics framework for the communication of the specific stereotaxic location of biorepository resources remains largely overlooked, despite the crucial importance of this information (Murray et al., 2025; Quinones-Hinojosa et al., 2024). The lack of spatial information for BSPs results in several barriers which limit the reproducibility of tissue-based studies, restrict interdisciplinary integration with neuroimaging, and lead to the underutilization of valuable disease relevant tissue samples.

Despite efforts including consensus guidelines for the assessment of neurological disease (Hyman et al., 2012; Montine et al., 2016) and pre-harmonization for data aggregation (Besser et al., 2018), there remains substantial variability in the anatomical landmarks used to locate sample regions and the sampling precision across centers remains unclear. For example, a survey of neuropathology cores in the Alzheimer’s Disease Research Centers found relatively high nominal consistency for the middle frontal gyrus (MFG) sampling, but also cautioned that landmark use remained heterogenous and that current approaches may be insufficient to ensure anatomical sampling consistency across centers (Vizcarra, Teich, Dugger, Gutman, & Alzheimer’s Disease Research Center Digital Pathology Working, 2023). Additionally, this survey found that typical samples for MFG were taken from coronal slices less than 5 millimeters thick, yet this gyrus can extend more than 10 centimeters along the anterior-posterior axis. The MFG contains multiple distinct cortical areas (Glasser et al., 2016) and along the MFG, studies have found systematic heterogeneity in patterns of connectivity (Jung, Lambon Ralph, & Jackson, 2022), cytoarchitecture (Bruno, Lothmann, Bludau, Mohlberg, & Amunts, 2024), gene expression (Wong et al., 2018), receptor architecture (Goulas et al., 2021), and disease-relevance (Haroutunian, Katsel, & Schmeidler, 2009). If the same sampled region across centers contains distinct populations of neurons with differing susceptibility to neurological disease, this could manifest as failures to replicate reliable results from other centers and artifactual variability in datasets which aggregate results across centers. These issues are likely to become increasingly relevant with the development of digital neuropathology tools and platforms for multi-institution whole slide digital imaging (Flanagan et al., 2025; Rosado et al., 2025).

Despite the inherently interdisciplinary nature of neurodegenerative disease research and the prevalence of interdisciplinary collaborations in the field, few studies robustly link neuropathology and antemortem neuroimaging. Even processes for linking neuropathology and *ex vivo* neuroimaging remain relatively rare and often require the development of elaborate infrastructure (Athalye et al., 2025; Faigle et al., 2023). In our experience, research studies utilizing human brain tissue rarely request samples from many regions known to be disease-relevant from neuroimaging studies. The neuropathologic assessment of brain tissue is the ground truth for many neuroimaging methods, yet often researchers will benchmark their results against other metrics. The lack of correspondence between the neuroscience terminology used by neuroimagers and the gross or microanatomical parcel names used by biorepositories complicates the identification of appropriate tissue resources. Conversely, neuropathologists and biomolecular scientists may be unfamiliar with the structure-function relationships for human brain regions which have been established through cognitive neuroscience and neuropsychology. With the rapid increase in the number of deeply phenotyped research participants who become brain donors through open science consortia such as the Alzheimer’s Disease Neuroimaging Initiative (ADNI), National Alzheimer’s Coordinating Center (NACC) Uniform Data Set (UDS), Standardized Centralized Alzheimer’s & Related Dementias Neuroimaging (SCAN), Consortium for Clarity in ADRD Research Through Imaging (CLARiTI), the Dominantly Inherited Alzheimer Network (DIAN), and Longitudinal Early-onset Alzheimer’s Disease Study (LEADS), there are also increasing research and clinical benefits to establishing widely adoptable methods for linking *in vivo* with *ex vivo* data.

Honoring the intent of brain tissue donors requires the efficient use of their tissue to address scientific questions, yet many tissue samples which are relevant to current research directions are rarely if ever requested. This issue is particularly salient in neurodegenerative disorders such as Alzheimer’s disease where the gold standard for etiological diagnosis, biomarker validation, and topographical distribution of disease processes is the assessment of postmortem brain tissue (DeTure & Dickson, 2019; Thal, Poesen, Vandenberghe, & De Meyer, 2025; VandeVrede et al., 2025). However, many brain tissue requests are for a small set of regions (e.g. “hippocampus and dorsolateral prefrontal cortex”), leaving most routine samples underutilized. Since the disease-relevance of these under-requested regions is primarily based on neuroimaging and clinical findings (Montine et al., 2012), there remains untapped potential for future interdisciplinary research. One factor in the underutilization of these disease-relevant tissues is the absence of a straightforward method for assessing the correspondence of neuropathological samples to the regions reported in the antemortem neuroimaging literature.

In this article, we describe a pipeline that biorepositories can use to create a Spatial Atlas Mapping the Protocol Locations of Ex vivo Samples (SAMPLES), which seeks to address these challenges and advance brain biorepository practices. Building on our previous work (Webster et al., 2021), we use a neuroimaging-based framework to create a normative digital atlas of a BSP on an advanced MRI brain template, resulting in a spatially and neuroanatomically explicit digital reference for BSP-defined samples.

We then illustrate the pipeline by creating a digital atlas for the fixed-tissue BSP developed by the University of Washington’s BioRepository and Integrated Neuropathology (BRaIN) Laboratory (Latimer et al., 2023), one of the most extensive sampling protocols in the field (Ellenbogen, 2024). Fixed tissue (e.g. Formalin-Fixed Paraffin Embedded (FFPE)) samples are nearly universally collected by biorepositories and provide the basis for neuropathological diagnosis (Shepherd, Alvendia, & Halliday, 2019). While most advanced biorepositories typically preserve around twenty fixed-tissue samples for subsequent histological analyses (Clement et al., 2021; Vizcarra et al., 2023), the BRaIN Lab BSP employs a modular gross sampling framework which includes standardized acquisition of 90+ samples (Supplementary Figure 1), covering a broad spectrum of functionally and neuropathologically significant brain regions (Latimer et al., 2023). These brain samples, along with extensive characterization including *ex vivo* neuroimaging, state-of-the-art preservation and staining techniques, and digital whole-slide imaging, provide an exceptionally high-quality resource for neurodegenerative disease research and neuroscience more broadly. Using this well-described protocol to illustrate the pipeline creates UW BRaIN SAMPLES, a precise spatial representation of the BRaIN Lab BSP.

**Figure 1.**
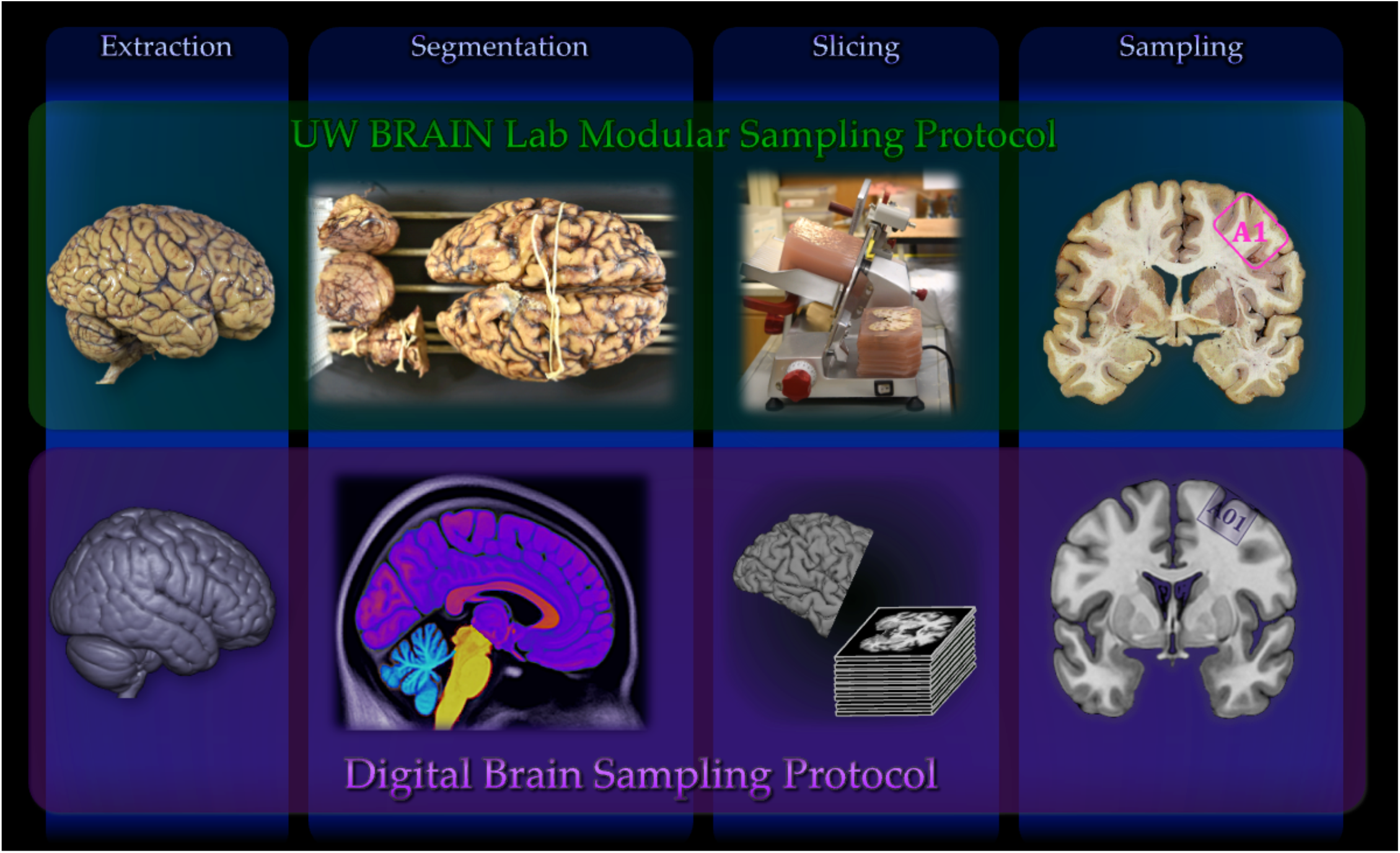
The Digital Brain Sampling Protocol. The Digital Brain Sampling Protocol is a metrically precise in-silico encoding the brain sampling protocol procedures for brain extraction, tissue segmentation, slicing, and sampling of all routine tissue samples of the protocol.

Our approach uses standard neuroinformatics conventions to create a 3D representation of a BSP through a metrically precise (real world units) digital instantiation of the procedures for brain extraction, segmentation, slicing, and sampling of the protocol. SAMPLES are normative references that can guide precision neuropathological sampling, facilitate the tissue request process, generate scientific visualizations, and provide a bridge between neuropathology and neuroimaging, in turn facilitating greater interdisciplinary insight from donated brain tissue.

## Materials and Equipment

Software, reference data, and equipment used to develop SAMPLES are listed in Table 1. The pipeline relies entirely on freely available analysis tools and one cloud-based vector-design application; no specialized hardware is required.

**Table 1.**
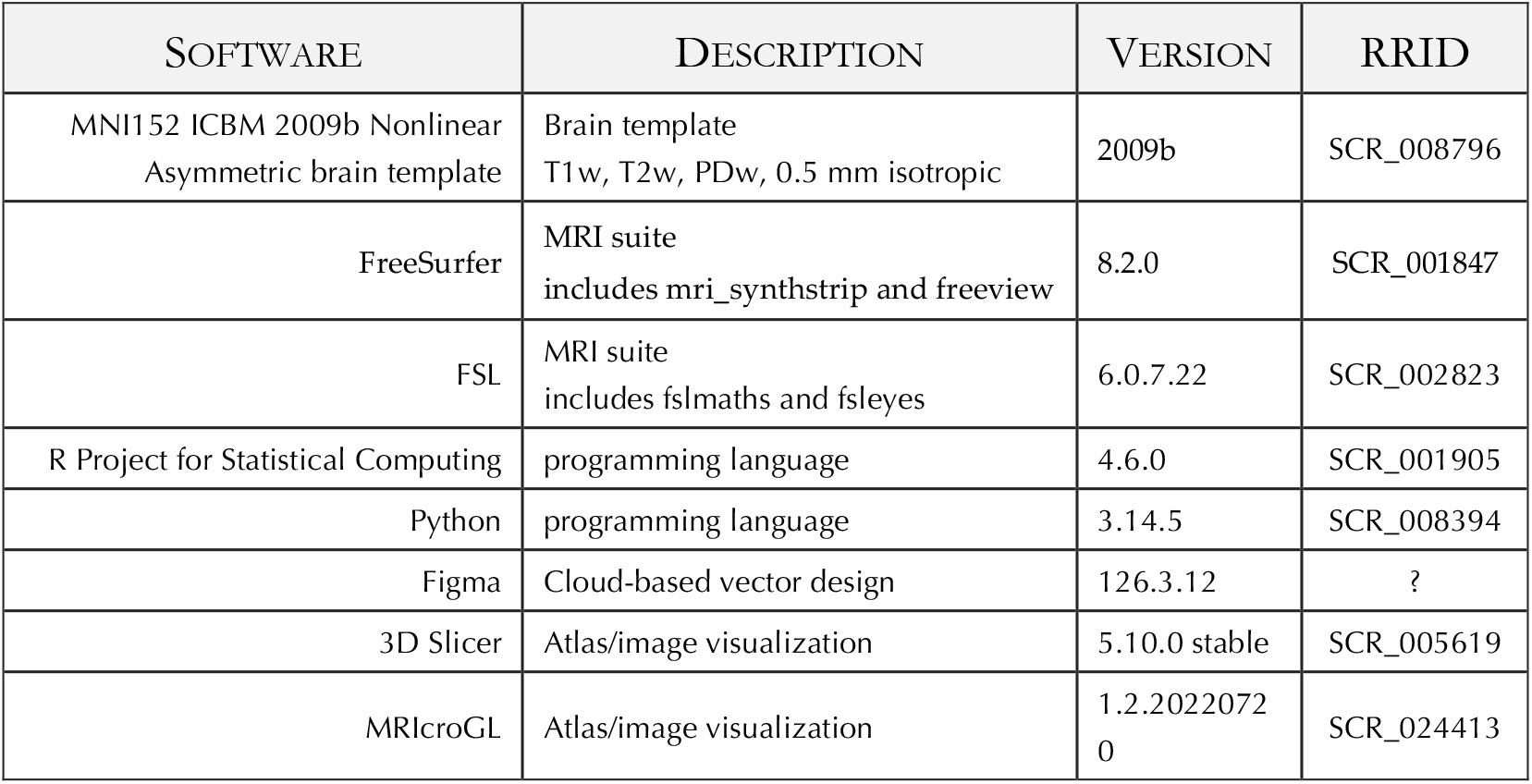
Resources used in the SAMPLES pipeline.

| SOFTWARE | DESCRIPTION | VERSION | RRID |
| --- | --- | --- | --- |
| MNI152 ICBM 2009b Nonlinear<br>Asymmetric brain template | Brain template<br>T1w, T2w, PDw, 0.5 mm isotropic | 2009b | SCR_008796 |
| FreeSurfer | MRI suite<br>includes mri_synthstrip and freeview | 8.2.0 | SCR_001847 |
| FSL | MRI suite<br>includes fslmaths and fsleyes | 6.0.7.22 | SCR_002823 |
| R Project for Statistical Computing | programming language | 4.6.0 | SCR_001905 |
| Python | programming language | 3.14.5 | SCR_008394 |
| Figma | Cloud-based vector design | 126.3.12 | ? |
| 3D Slicer | Atlas/image visualization | 5.10.0 stable | SCR_005619 |
| MRICroGL | Atlas/image visualization | 1.2.20220720 | SCR_024413 |

## Methods

A Spatial Atlas Mapping Protocol Locations of Ex vivo Samples (SAMPLES) is based on a high-resolution digital brain template and is generated by following the digital brain sampling protocol (dBSP) pipeline. By computationally specifying the details for brain extraction, segmentation, slicing, and sampling (Figure 1), this pipeline captures the metric and neuroanatomical properties of samples collected according to a particular brain sampling protocol. SAMPLES uses standard neuroinformatics conventions which facilitate interfacing with openly available neuroimaging software. SAMPLES was developed using FMRIB Software Library (FSL; https://fsl.fmrib.ox.ac.uk/), FreeSurfer (http://surfer.nmr.mgh.harvard.edu/), R (https://www.r-project.org/), and python (https://www.python.org/). Italicized terms in this section indicate specific software tools or intermediate files used in the construction of SAMPLES.

The UW BRaIN SAMPLES models the UW BRaIN Laboratory’s collection and modular gross sampling protocol (Latimer et al., 2023) for FFPE tissue. The atlas contains all routinely collected samples (n=90), comprising five modules: routine diagnostic (22), brainstem (17), chronic traumatic encephalopathy (CTE) (14), traumatic brain injury (TBI) (9), and neuroimaging (28) (see Supplementary Materials, Figure 2).

**Figure 2.**
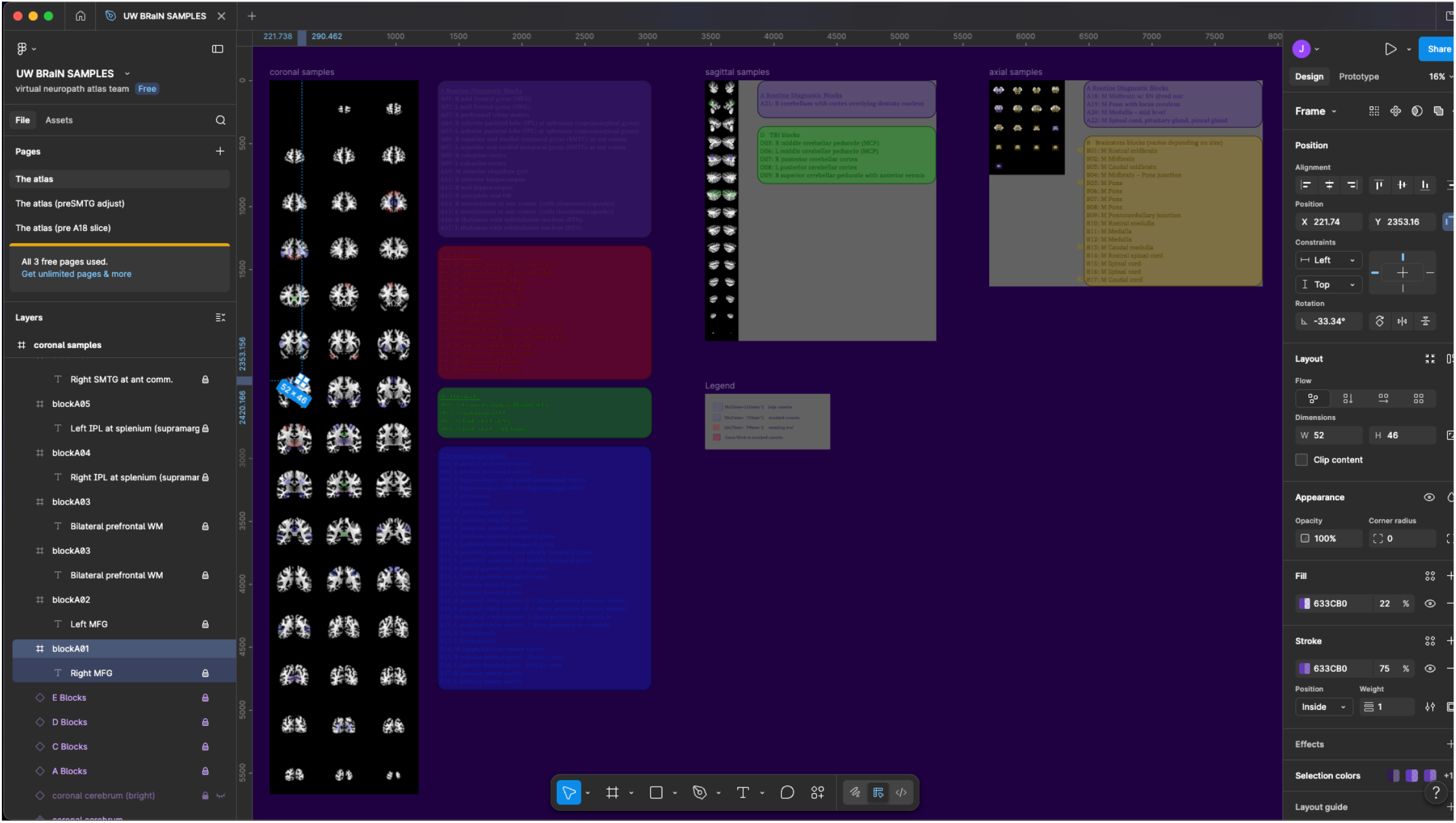
The Virtual Sampling Interface. Virtual neuropathology sampling was implemented in Figma, a cloud-based design tool with an intuitive and responsive interface. For each panel (cerebrum, coronal; cerebellum, sagittal; brainstem, axial), the corresponding samples names and descriptions were color coded according to sample module. Virtual sampling was performed by a neuropathologist and the lead tissue procurement technician. Samples taken with the sampling tool were indicated by dragging the labelled rectangle next to each corresponding sample name to the appropriate slice then adjusting the position and rotation. Samples excised by scalpel were indicated by dragging the labelled rectangle next to each corresponding sample name to the appropriate slice then drawing a closed-contour with the pen tool. Data on sample position were exported in Scalable Vector Graphics (svg) format.

### Brain Template

The landmarks for BSP sampling in practice are the gross anatomical features consistent across donor brains. However, the morphology, proportions, and distance of these features vary between neurotypical brains and are extensive with the cases of end-stage neurological disease that are the focus of many biorepositories. SAMPLES atlases are based on digital brain templates to explicitly encode the normative spatial and neuroanatomical specifications for a BSP. Created by computationally aligning and averaging structural magnetic resonance imaging (MRI) data from many individuals, digital brain templates represent consistent neuroanatomical features in a metrically specified common coordinate framework.

UW BRaIN SAMPLES is based on the Montreal Neurological Institute (MNI) 2009b International Consortium for Brain Mapping (ICBM) 152 Nonlinear Asymmetric Brain (*the MNI2009b brain*) (V. Fonov et al., 2011; V. S. Fonov, Evans, McKinstry, Almli, & Collins, 2009). *The MNI2009b brain* is available in three MR image contrasts (T1w, T2w, and PDw) with a resolution of .5 mm isotropic, eight times finer than typical research-grade structural MRI data and brain templates in most neuroimaging software packages.

### Brain Extraction

During brain extraction, brain tissues for further processing are isolated from non-brain structures (e.g. skull, dura, meninges, external vasculature). Including this stage in the digital BSP ensures the separation of all brain tissue from adjacent non-brain tissue for subsequent processing as the *brain-extracted volume*, ensuring that non-brain structures adjacent to sampled tissue are not labelled in the final atlas. This volume is created by removing (setting to zero) all non-brain tissue as specified in the BSP from the digital brain template.

The precision required to create *brain-extracted volume* for UW BRaIN SAMPLES required a semi-manual procedure. Briefly, *the MNI2009b brain* was duplicated, processed with FreeSurfer’s *mri_synthstrip* (Hoopes, Mora, Dalca, Fischl, & Hoffmann, 2022), thresholded to remove voxels containing primarily CSF, and manually edited in FreeSurfer’s *freeview* to remove non-brain structures and restore any omitted brain tissue (see Supplementary Methods).

### Brain Segmentation

During brain segmentation, anatomic segments of the brain (e.g. brainstem, cerebellum, and cerebrum) are physically separated before further processing. Including this stage in the digital BSP ensures accurate reflection later of BSP stages which differ for different brain segments and are particularly important for assigning tissue to correct samples at segmentation boundaries. For each segment, a cropped *volume* is created from the *brain extracted volume* which only includes the corresponding BSP-defined anatomical segment. This will likely require a semi-manual process since the BSP-specified cuts are likely to be inclined or curved relative to the voxel axes and are unlikely to precisely correspond to available automated anatomical segmentation boundaries.

In the UW BRaIN Laboratory BSP, the posterior fossa contents are separated by a plane from the base of the mammillary bodies to the base of the thalamus, and then the brainstem and cerebellum are separated laterally along the curvature of the pons through the cerebellar peduncle and along the anterior cerebellum through the cerebellar peduncles and medullary vermis. A semi-automated approach combining computational masking and manual refinement to ensure precise correspondence with this segmentation protocol (see Supplementary Materials for more detail). First, the *brainstem volume* was initialized from the *brain-extracted volume*, multiplied by a binary mask for the bounding box of the brainstem (i.e. setting voxels outside the bounding box to zero), and refined in freeview to remove voxels corresponding to the cerebrum and cerebellum. The *brainstem mask* is the binarized *brainstem volume*. Next, the *cerebellum volume* was initialized from the *brain extracted volume*, multiplied by the inverted bounding box of the cerebellum, multiplied with the inverted *brainstem mask*, and iteratively refined in *freeview* to remove voxels corresponding to the cerebrum. The *cerebellum mask* is the binarized *cerebellum volume*. Finally, the *cerebrum volume* was initialized from the *brain extracted volume*, multiplied with the inverted *brainstem mask*, then multiplied with the inverted *cerebellum mask*. The *cerebrum mask* is the binarized *cerebrum volume*.

### Brain Slicing

Brain slicing creates a series of slices according to BSP specifications for slice thickness and angle relative to the anatomical dimensions of the brain segments. Including this stage in the digital BSP ensures the correct mapping between slice order, orientation, face, and thickness for subsequent stages. For each segment, if the axis along which slices are taken does not correspond to a voxel dimension, the data is rigidly rotated to align the axis to nearest voxel dimension and resampled to create a rotated segment *volume*. Then 2D *brain slice images* are created as png files using custom R code along the corresponding voxel dimension at indices increased as indicated by the BSP-specified slice thickness.

For the UW BRaIN Laboratory fixed-tissue protocol, the relevant details to capture during this stage were the slice orientation, initial brainstem slice, and slice thickness.

In *the MNI2009b brain*, coronal slices of the cerebrum and sagittal slices of the cerebellum correspond to planes identified by indexes along dimensions of volume. While the remaining vertical dimension of the volume corresponds to the axial dimension of the cerebrum, the brainstem is sliced rostral to caudal, along the long axis of the brainstem, which is pitched forward relative to vertical within the volume. Thus, an affine transformation was applied to the *brainstem volume* that performs the minimum rotation around the volume centroid to pitch the first principal component axis of the central sagittal slice to align with the vertical axis to create the *vertical brainstem volume*.

The initial brainstem slice is cut parallel to the anatomically determined segmentation cut to ensure that sample A18, which is crucial for routine neuropathological diagnosis, has uniform thickness and includes the specified structures (midbrain with substantia nigra and red nucleus). As these cutting planes are slightly anteriorly tilted relative to the horizontal plane of *the vertical brainstem volume*, the segmentation cut was computationally operationalized by lowering a plane until it contacted tissue then pivoting forward to the segmentation cut. Using a coordinate frame for the plane and corresponding normal vector centered on the pivot, the cutting plane for the initial brainstem slice was determined. The *initial brainstem slice image* was created from the intersected voxels. Voxels corresponding to sample A18 were identified computationally by positioning rectangular mask matched to the sampling tool dimensions on the segmentation plane and ‘cutting’ through the slice, then copying the A18 atlas label into these voxels into the *brainstem sample volume*. The initial brainstem slice was then removed from the *vertical brainstem volume*.

Finally, since the MNI2009b brain has .5 mm isotropic voxels, the *brain slice images* were then created as png files every 8 voxels (4 mm) from anterior to posterior for *cerebrum volume*, medial to lateral sagittally for the *cerebellum volume*, and rostral to caudal for the *vertical brainstem volume*.

### Virtual Sampling Interface

BSPs specify how brain slices are systematically oriented and arranged for sampling, how many samples are taken, what the samples should include, and the spatial dimensions of the samples. Including this stage in the digital BSP ensures samples are in the correct hemisphere and that the slice thickness is added back along the correct direction when the digital samples are later converted to 3D. A virtual neuropathology sampling environment (Figure 2) was developed in Figma, a cloud-based design tool that facilitates real-time remote collaboration. Figma provides an intuitive online graphical user interface (GUI) that facilitates precise and efficient interaction with keyboard, mouse, trackpad, or touchscreen. Design elements can be locked to prevent accidental changes. Data are exportable in scalable-vector graphics (SVG) which is a lossless representation that ensures metric accuracy. A frame was created for each segment with *brain slice images* on the left, text for each sample on the right, and a sample template for each uniform tissue block next to the text for the corresponding sample text. The upper left of each frame has a coordinate of (0,0) and slices are tiled to facilitate the mathematical extraction of pixel coordinates.

For the UW BRaIN Laboratory protocol, *brain slices images* were laid out on three frames in Figma for the coronal cerebrum slices, sagittal cerebellum slices, and off-axial brainstem slices. Text boxes for each sample module were displayed on the right of each frame which contain the name, hemisphere (e.g. L (left), R (right), M (medial), or B (bilateral)), and description (e.g. “E11: L fusiform / inferior temporal gyrus”) of each corresponding sample. A rectangular frame for each sample in the BSP was placed next to the corresponding line in the text box. Each frame shows the corresponding sample ID (e.g. E11) and displays an associated text description (e.g. L fusiform / inferior temporal gyrus) upon mouseover. For actual samples taken with the sampling tool, sample frames showed a 52×46 pixels rectangular border while those corresponding to actual samples taken by scalpel excision had no visible border.

Colors were used consistently for each module’s text box, sample ID, and frame border. All elements except the sample frames were locked to prevent accidental modification.

### Virtual Neuropathology Sampling

Sampling is typically guided by the laterality, position, orientation, and anatomical targets specified by the BSP and carried out by trained experts to improve consistency across cases. Including this stage in the digital BSP explicitly establishes the normative targets for samples according to the BSP. Selecting one of the sample frames jumps to the corresponding frame element in the left navigation panel and mousing over a rectangle highlights the corresponding frame element. Conversely, clicking on a frame element in the left navigation panel selects the corresponding sample frame and provides crosshair for rapid identification. The sample frames for tissue blocks are dragged to the appropriate slice and precisely positioned by translation and rotation, aided by smooth and rapid zooming to optimize the placement. For virtually excised samples, the pen tool is used to create a closed contour stored that is stored as a vector object within the frame.

For the UW BRaIN Laboratory protocol, a neuropathologist and the lead tissue procurement technician used the online interface to conduct the virtual neuropathology sampling procedure, indicating each tissue block and excision (Figure 3).

**Figure 3.**
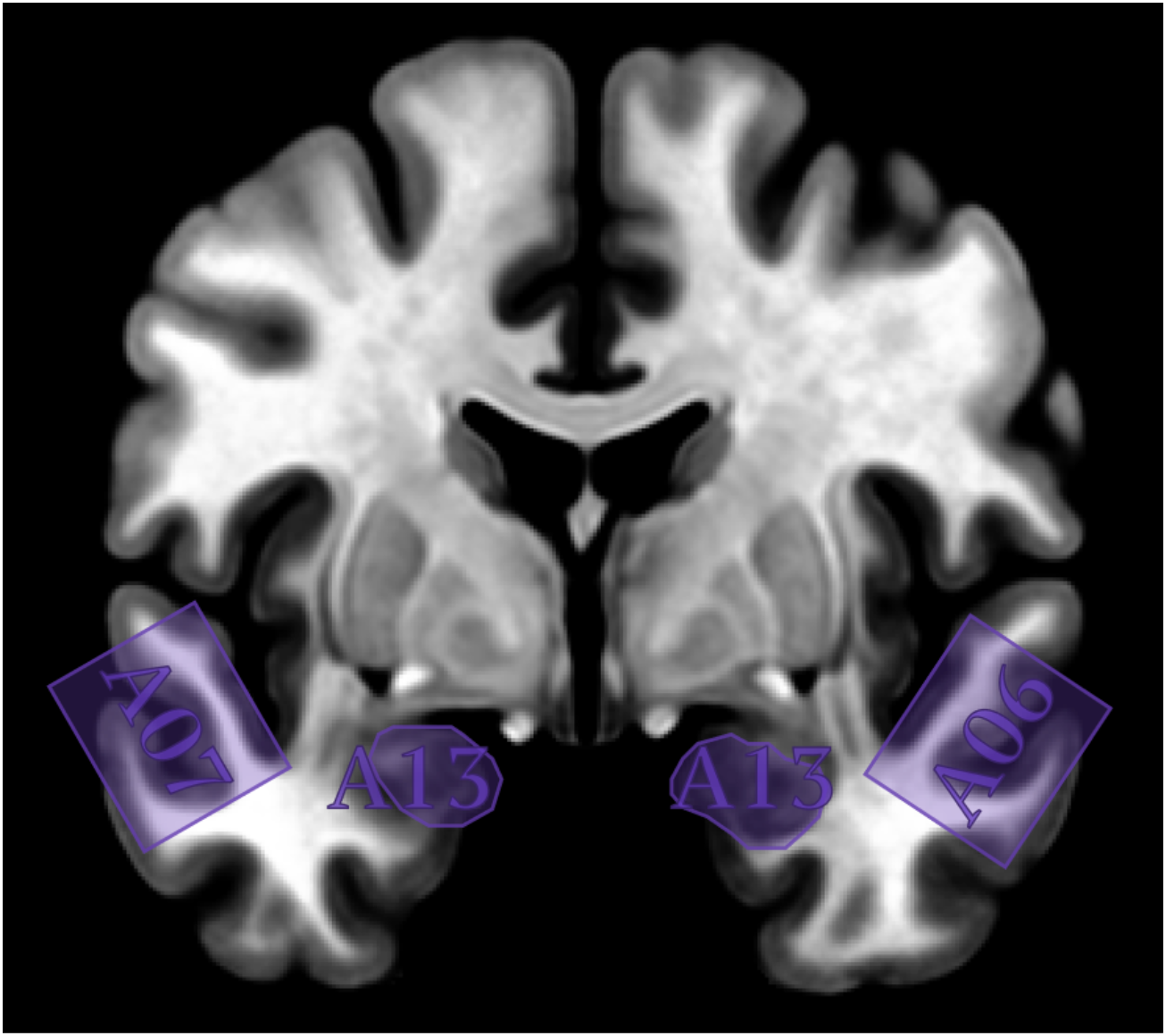
Example brain slice from the Virtual Sampling Interface. A coronal slice of the cerebrum showing samples taken with the sampling tool (A06, A07) and by excision (A13).

Once all samples had been placed in the consensus locations, data were exported in SVG format for cerebrum, cerebellum, and brainstem frames.

### Digital Atlas Creation

Digital atlas creation transforms the slice-based sample placements produced during virtual sampling into a 3D label atlas that encodes them in template space. This stage must convert 2D sample geometries into volumetric extents consistent with BSP-defined slice thickness and sampling tool geometry, enforce component-specific tissue constraints (e.g., restricting labels to the appropriate segmented volume), and integrate outputs across components into a single atlas representation with stable label identifiers and metadata. Accurate coordinate transforms, slice-to-volume mapping, and thickness extrusion are particularly important because errors at this stage systematically distort the spatial extent and location of the resulting targets.

The files constituting SAMPLES are generated using custom python code (https://www.python.org/). Information regarding the BSP is read in from a spreadsheet which included the index, ID, hemisphere, type, and module of each sample (if applicable). Spatial information from the virtual neuropathology sampling is read in from the SVG files obtained from Figma.

For UW BRaIN SAMPLES, the *sample volumes* for the *cerebrum, cerebellum*, and *brainstem* were initialized as zero-filled matrices with dimensions matching the *cerebrum volume, cerebellum volume*, and *vertical brainstem* volume. For each sample, the 2D coordinates from the SVG sample locations were converted to 3D right, anterior, superior (RAS) coordinates in the corresponding volume from the brain segmentation. These coordinates were then connected as closed paths, converted to filled polygons, and replicated through 8 slices along the orthogonal dimension in the direction specified by the BSP (in this case anterior for cerebrum samples, superior for brainstem samples, and lateral for cerebellum samples. All voxels contained within this 3D volume were then filled with the index value of the corresponding sample. To constrain the sample indices in the atlas to the brain tissue in corresponding segments, the indices were multiplied by their respective binary segmentation mask. The *brainstem sample volume* was then created from the *vertical brainstem sample volume* by applying the inverse of the affine used to create the *vertical brainstem volume* (returning the brainstem samples to MNI2009b space).

The *UW BRaIN SAMPLES volume* was initialized as a zero-filled volume with dimensions corresponding to the *MNI2009b brain*, the indices from the *cerebrum, cerebellum*, and *brainstem sample volumes* were transferred into the *UW BRaIN SAMPLES volume*, and the atlas was saved in compressed NIFTI format with intention set to “Labels”.

Additional files were generated for the compatibility of SAMPLES with commonly used neuroimaging software. A FreeSurfer lookup table was created to link the sample indices, names, and RGB values for visualization in FreeSurfer’s *freeview* as well as other software which accept this standard such as FSL’s *fsleyes* and 3D Slicer. Another lookup table with 256 8bit unsigned integers for red, green, and blue values and a label file with sample indices and names was generated for visualization in MRIcroGL.

### UW BRaIN SAMPLES Verification

The accuracy of atlas labels was determined through systematic inspection. The MNI2009b brain, UW BRaIN SAMPLES, and Destrieux anatomical atlas (Destrieux, Fischl, Dale, & Halgren, 2010) were loaded in *freeview*. Initial visual inspection was conducted for each sample by selecting it in the Lookup Table panel then traversing the coronal, sagittal, and axial views to evaluate sample placement. First, the hemisphere label in the sample description was compared to the Coordinate Annotation. The relative location within the brain was assessed based on guidance from the BSP. The specific placement of the sample was compared to the placement in Figma virtual neuropathology sampling interface. The direction and extent for the thickness of the sample was assessed, in this case eight voxels from the slice number where the sample was placed in virtual sampling interface along the dimension specified by BSP. Any anatomical structures directly mentioned in the sample description were checked against the labels provided by the Destrieux atlas in the information panel. Structures not included in the Destrieux atlas, such as the locus coeruleus or red nucleus, were visually identified through adjusting the image contrast of the MNI2009b brain and assessed for inclusion in the corresponding sample.

## Results

Here, we introduce the digital BSP pipeline, a process that biorepositories can adapt to generate a Spatial Atlas for Mapping Protocol Locations of Ex vivo Samples (SAMPLES).

We demonstrate an instance of such a process for a leading biorepository for Alzheimer’s disease and related dementias. UW BRaIN SAMPLES represents the UW BRaIN lab BSP mapped onto the MNI2009b brain. The atlas consists of a 3D NIFTI volume of 16-bit integer values representing sample label IDs as well as the files linking these label IDs to sample information and colors for display for both FreeSurfer (Figure 4) and MRIcroGL (Figure 5).

**Figure 4.**
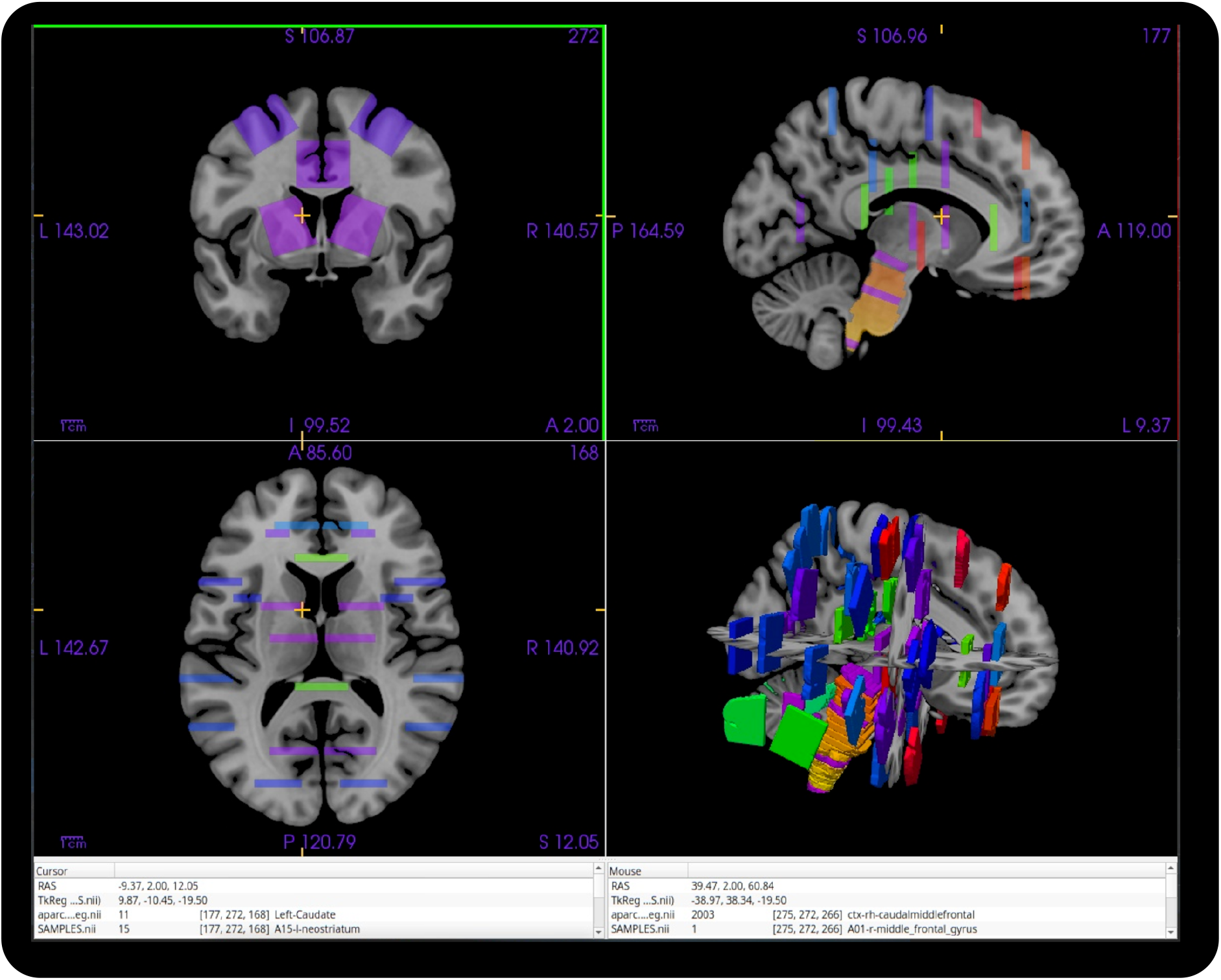
Interactive visualization of a digital brain sample protocol atlas in FreeSurfer’s *freeview* interface. The University of Washington BioRepository and Integrated Neuropathology Spatial Atlas Mapping Protocol Locations for Ex vivo Samples (UW BRaIN SAMPLES) is shown. Sample color corresponds to module (purple: diagnostic, yellow: brainstem, red: CTE, green: TBI, blue: neuroimaging). Sample information is shown in the information window for the cursor position (left pane) and mouse position (right pane).

**Figure 5.**
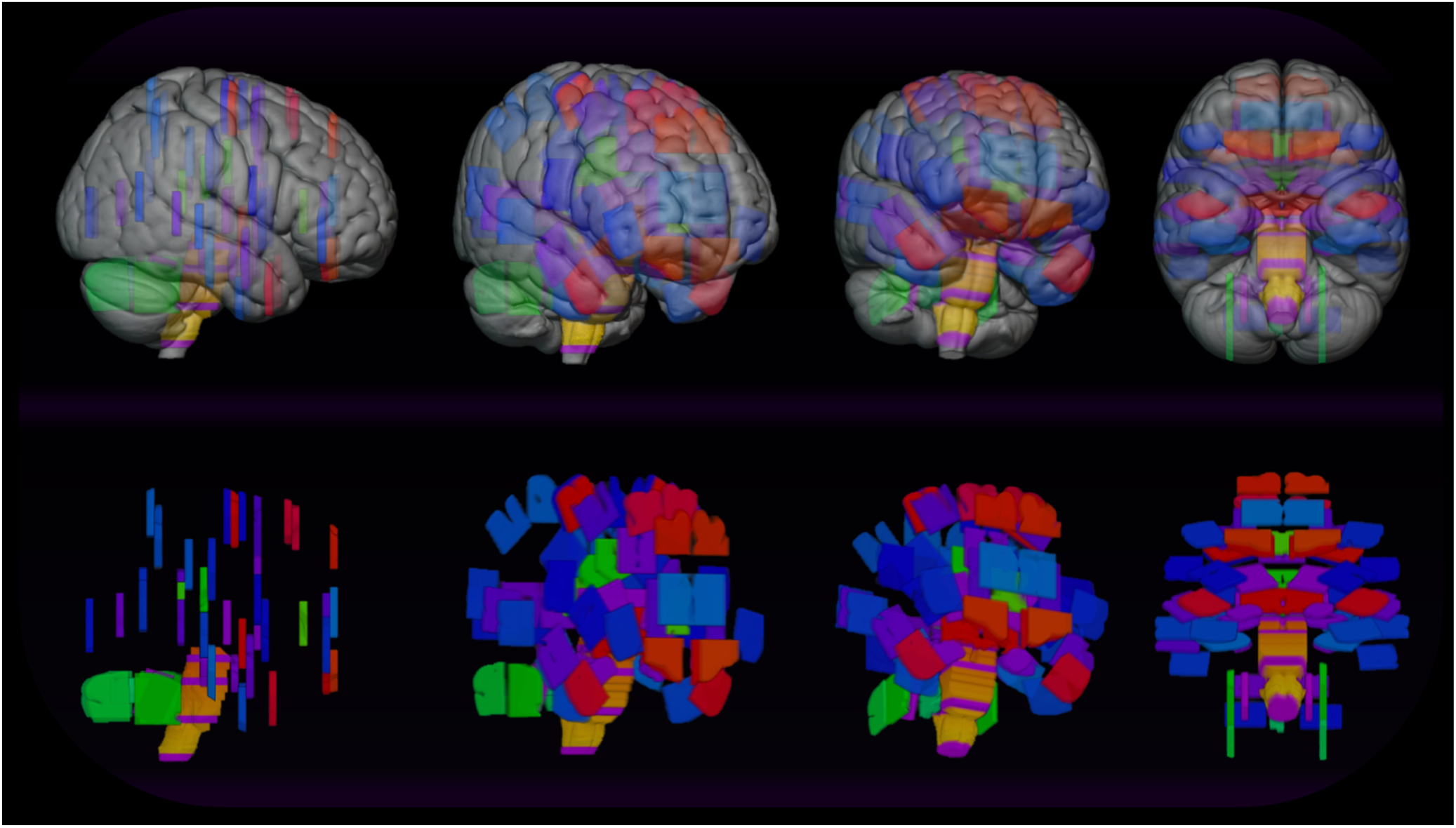
Interactive 3D visualization of the position of routinely collected fixed tissue samples displayed in MRIcroGL. The University of Washington BioRepository and Integrated Neuropathology Spatial Atlas Mapping Protocol Locations for Ex vivo Samples (UW BRaIN SAMPLES) is shown. Sample color corresponds to module (purple: diagnostic, yellow: brainstem, red: CTE, green: TBI, blue: neuroimaging).

For each virtually positioned sample, anatomic correspondence was verified with sample placement in the virtual neuropathology sampling interface as well as guidance from the BSP (e.g. “on the same slice as the anterior commissure”, “two slices posterior to block E27”). Samples whose descriptions mention specific neuroanatomical structures were confirmed to contain the corresponding Destrieux atlas labels. For those with corresponding labels in the Destrieux atlas, the overlap of those labels with sample was established in *freeview*. For example, in Figure 6, sample A01 was positioned in the right hemisphere and encompassed the middle frontal gyrus, with the corresponding Destrieux atlas label (*ctx_G_front_middle*) confirming its anatomical accuracy.

**Figure 6.**
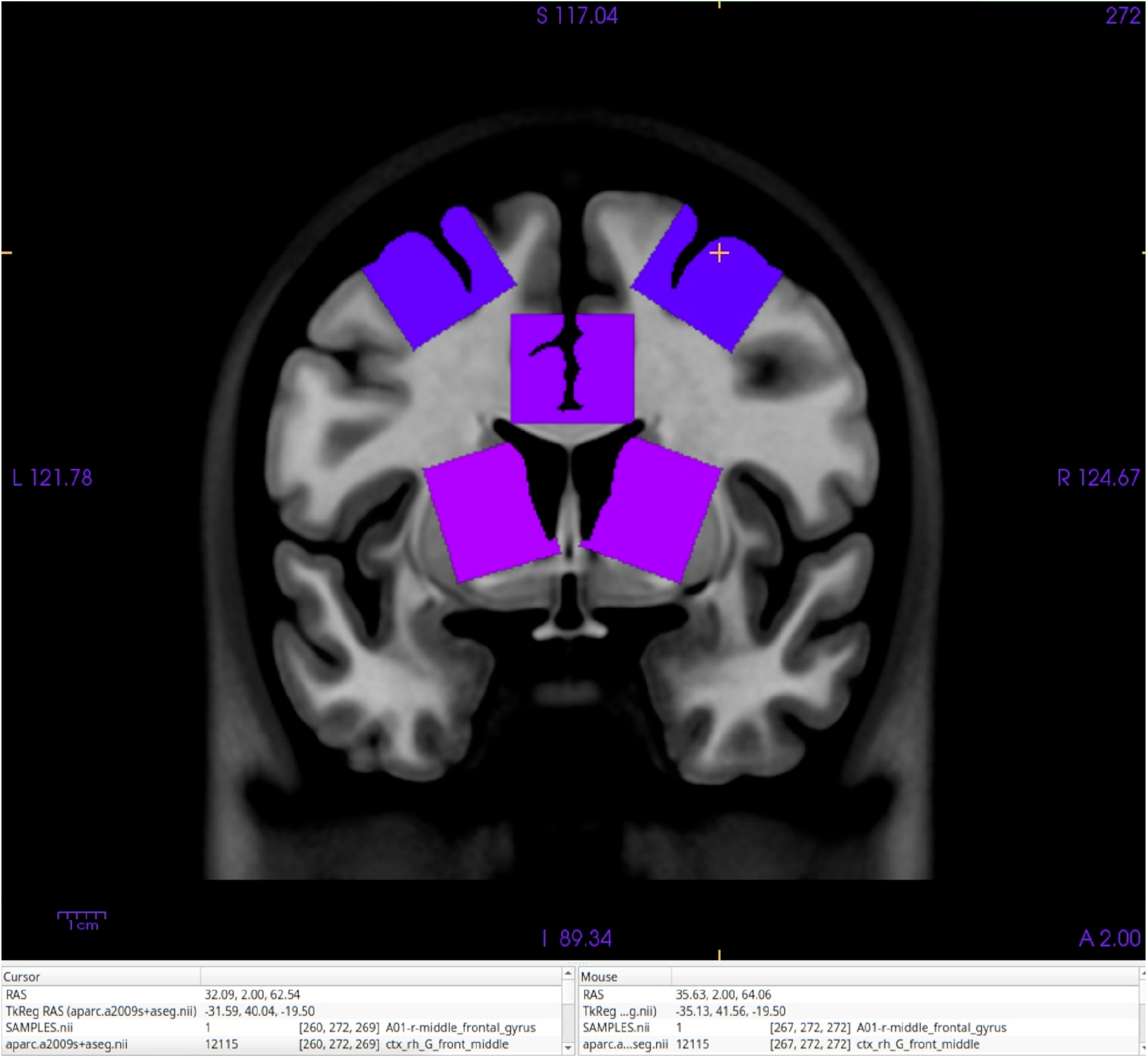
The verification process using FreeSurfer’s *freeview* is shown for sample A01: Right Middle Frontal Gyrus. The Information Pannel at the bottom of the figure shows the cursor (gold cross) is on A01-r-middle_frontal_gyrus. The ‘r’ indicates the sample is in the right hemisphere, matching the “R” in the Coordinate Annotation (blue text) on the left of the screen. The sample is in the frontal lobe, near the anterior commissure (not visible from this slice), contains the middle frontal gyrus and sulcus, and is rotated to align with the pial surface. In the Cursor Information panel at the bottom of the screen, the Destrieux (aparc.a2009s+aseg) label is verified to be ctx_rh_G_front_middle (the right Middle Frontal Gyrus).

Subcortical structures without corresponding labels in the Destrieux atlas were first visually identified in *fsleyes* and confirmed to be with the appropriate samples, as exemplified by sample A19 (Figure 7), which contained the pons and locus coeruleus. These results confirmed that the UW BRaIN SAMPLES provides a spatially accurate representation of the brain sampling protocol, as well as the ease of use in various neuroimaging software packages.

**Figure 7.**
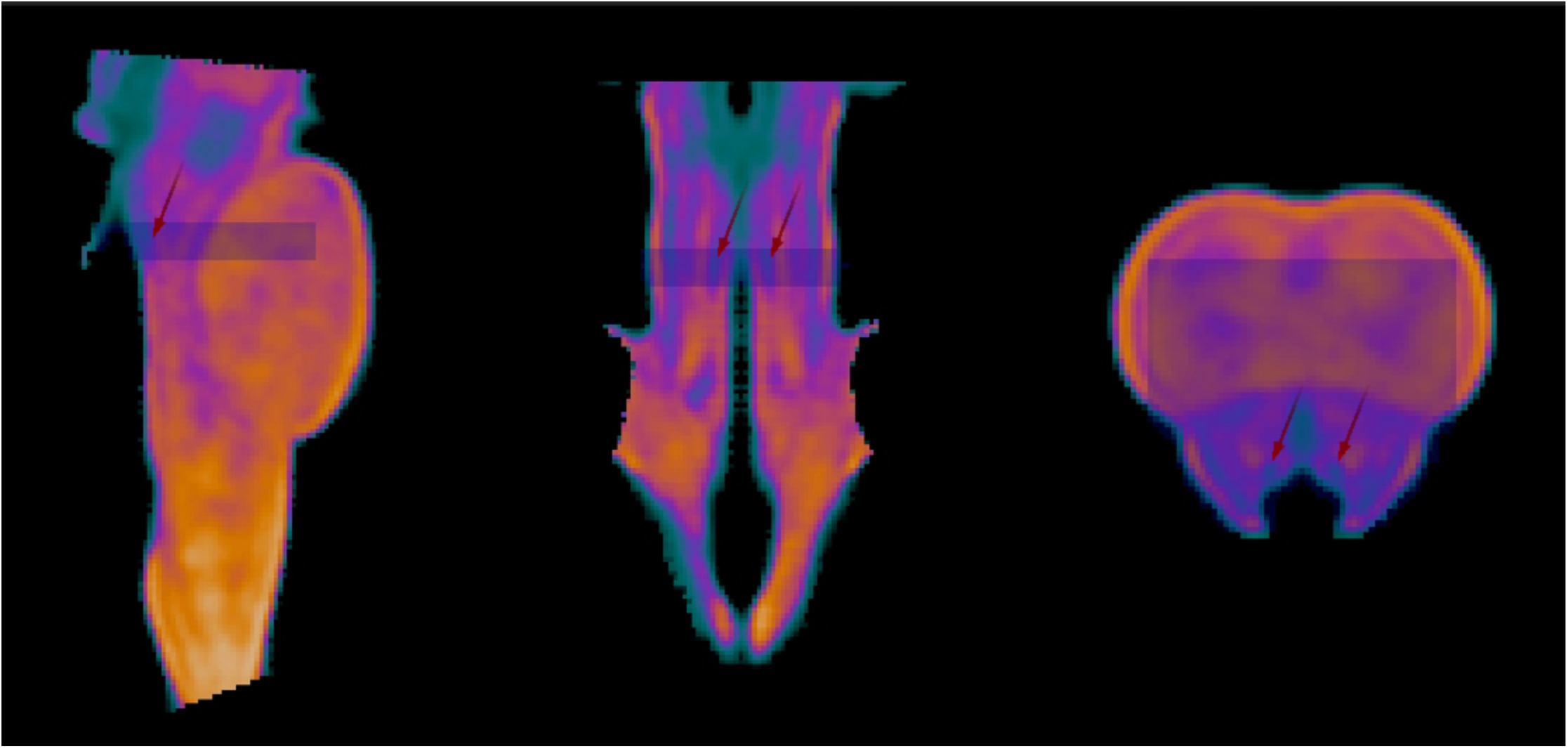
Verification process using FSL’s *fsleyes* for sample A19: Medial Pons with Locus Coeruleus. The sample contains the pons and locus coeruleus (red arrows).

## Discussion

Encoding brain biorepository tissue resources as normative digital atlases on a modern neuroimaging template provides an explicit 3D representation of tissue samples in an established common coordinate framework. Spatial Atlas for Mapping Protocol Locations of Ex vivo Samples (SAMPLES) atlases are created through the digital brain sampling protocol pipeline that digitally implements brain extraction, segmentation, slicing, and sampling on a modern neuroimaging template. Each stage of the pipeline consists of capturing site- and protocol-specific details that result in metric and neuroanatomical correspondence with the BSP.

Our approach represents *intended* sample locations as quantitative, computation-ready volumetric labels on a brain template with maximally typical anatomy. While most biorepositories have specific standard operating procedures (SOPs) to consistently target the same neuroanatomical location for each sample across donors, our estimates suggest substantial variability remains in both the precision and accuracy of sampling from a combination of avoidable and unavoidable sources (Webster et al., 2021). Even taken with maximal location precision, some differences in tissue composition are likely due to individual differences in cortical folding, cortical area sizes, subcortical structure volumes, or the effects of neurobiological disease. A related study used an MRI brain template to create labels for the maximum volume of tissue from which pathologists might sample for eight sample locations (Raman et al., 2016), which reflects sampling *precision*. Although we also use imaging methods to track sample-anatomic correspondences, our approach is distinct in that we provide normative atlases of the *target* sample locations, which could be used to assess sampling *accuracy*.

### UW BRaIN SAMPLES

This article illustrates the atlas generation process by adapting the pipeline to create UW BRaIN SAMPLES, an atlas of the routinely collected fixed-tissue samples from the UW BRaIN Lab BSP. This is a leading tissue biorepository for Alzheimer’s diseases and related degenerative conditions embedded in a longstanding AD Research Center that is part of the nationwide network of ADRCs funded by the US National Institutes of Health, that samples the brain extensively and that shares samples and data prolifically. Planned extensions to the UW BRaIN SAMPLES include the addition of the frozen-tissue brain sampling protocol, incorporating intermittently collected samples for specific projects or collaborators, and creating historical atlases which represent prior versions of the BSP.

### Implications for Tissue Requesters

The lack available spatial information and included brain regions for brain biorepository resources can delay the identification of research relevant samples and complicate the comparison of results from tissues across biorepositories (LaBaer et al., 2018). For researchers requesting postmortem tissue, SAMPLES reduces uncertainty in sample location, supports resource discovery, and facilitates interdisciplinary research. Unlike labels for comparatively much larger Brodmann Areas or cortical gyri, the volumetric labels from SAMPLES reflect the dimensions of the corresponding tissue blocks, where the samples are taken within large parcels, and all the structures which each sample typically contains. During the tissue request process, SAMPLES provides an explicit, interactive map of the brain tissue available for request. These maps can be compared visually or computationally to identify biorepositories whose samples overlap, which could reduce variability when aggregating data across sites by excluding data from samples which likely contain different neural populations. SAMPLES can also lower the barrier to interdisciplinary research by facilitating the identification of tissue samples corresponding to research-relevant regions from the literature of other disciplines.

### Implications for Biorepositories

A unified framework for characterizing the spatial properties of tissue samples is a crucial addition to the ongoing efforts to standardize biorepository procedures and would further enhance harmonization efforts for cross-site data integration to enable scalable and reproducible discoveries (Murray et al., 2025). The SAMPLES pipeline is designed to generate an explicit spatial reference for the fixed-tissue resources preserved and made available for scientific research by brain biorepositories.

SAMPLES are a flexible computable resource which could enhance biorepository workflows including managing internal resources, training neuropathology technicians, guiding neuropathology sampling, implementing quality control measures, referencing available tissue samples, and augmenting tissue request meetings.

The ability to dynamically explore an explicit spatial atlas of the BSP would also facilitate communication between biorepositories and stakeholders for the future extension or revision of the BSP such as the addition of new samples based on neuroscience properties reflected in existing digital atlases, adjusting sample positioning to include structures of interest, or even redefining optimal sample targets as a specific distribution of specified brain structures.

Currently, there is little standardization in the number, location, or dimensions of the samples taken across brain biorepositories. Even when samples across biorepositories were derived from the same guidelines with ongoing harmonization efforts, anatomical correspondence is not assured and may differ substantially and systematically across institutions (Lucot et al., 2023; Vizcarra et al., 2023). Where possible, the approach and software described here anticipate these differences, however, ensuring accurate SAMPLES creation for a new BSP requires that each step of the dBSP pipeline captures relevant site and protocol specific properties including where segmentation cuts are made, how segments are oriented during slicing, slice thickness, how slices are arranged for sampling, and how the location of samples are selected. The utility and benefit of these SAMPLES atlases increase with increasingly widespread implementation across biorepositories.

Comparison of SAMPLES atlases across institutions could inform the interpretation of discrepant findings and enhance harmonization efforts. The dBSP pipeline brings sampling protocols into the common coordinate framework of a digital brain template, where the correspondence of sampling practices across institutions can be assessed through established neuroimaging approaches for interactive visualization and quantitative comparison.

The ability of researchers to easily identify the availability of disease relevant tissue resources could contribute to much greater utilization of samples which are rarely requested.

### Implications for Interdisciplinary Research

Recent estimates of the financial costs of the lack of reproducibility in preclinical research were estimated to in the tens of billions of dollars in the United States alone(Freedman, Cockburn, & Simcoe, 2015). One potential source for these failures of replication is variability in targeting the same brain areas, and hence cell populations, as the original studies. Researchers seeking to replicate prior work on studies using tissue from biorepositories with SAMPLES could computationally determine which biorepositories have sampling protocols that contain the same portions of brain tissue.

Basing SAMPLES in a neuroimaging framework provides a bridge for the scientific concepts and terminology between *ex vivo* and *in vivo* researchers. Neurodegenerative disease researchers with neuroimaging expertise may recognize the research value of under-requested tissues and have the software expertise to quickly incorporate a SAMPLES atlas into neuroimaging analysis pipelines. SAMPLES supports clinicopathologic workflows by computationally relating neuropathologic findings to premortem neuroimaging data. Supporting these novel research connections will require the expertise of biorepository members as well as molecular and cellular biologists.

Careful consideration should be given in interpreting results from applying any atlas to individual structural MRI. The MNI2009b template will differ from the anatomy of an individual, especially for older adults and those with neurodegenerative disease, and registration accuracy depends on both the algorithm and the anatomical characteristics of the individual brain, particularly in the presence of atrophy or structural deformation ((Dickie et al., 2017)). Moving beyond normative templates, we are exploring postmortem imaging modalities to estimate where samples were actually taken for each donor, allowing for ongoing assessments of precision and accuracy at the individual case level.

SAMPLES works with many freely available neuroimaging software packages that have highly customizable static and interactive visualizations for generating figures for the publication and presentation of scientific research.

### Future Directions

Ongoing efforts building on this work include developing a series of static and dynamic visualizations that serve as reference resources for biorepository workflows including interactive visualizations for tissue sample requests. To further facilitate resource discovery, a system is being developed to leverage digital neuroscience atlases from a range of fields to enable automated identification of samples containing tissue associated with nearly any neuroscience term.

In the longer term, we will use SAMPLES to develop virtual neuropathology training software and prospective sample guidance to further enhance the precision and accuracy of neuropathological sampling.

## Conclusion

The digital brain sampling atlas pipeline uses a neuroimaging framework to digitally instantiate procedures for brain extraction, segmentation, slicing, and sampling according to a specific BSP in order to accurately reflect the metric and neuroanatomical properties of the brain tissues being sampled. The resulting Spatial Atlas for Mapping Protocol Locations of Ex vivo Samples (SAMPLES) is a normative 3D representation of a particular BSP on a high-resolution MRI brain template.

We demonstrate the approach by applying the pipeline to generate UW BRaIN SAMPLES, a metrically accurate digital representation of a state-of-the-art, modular fixed-tissue protocol from a leading Alzheimer’s Disease Research Center.

By explicitly characterizing the spatial properties of samples in a BSP using an established common coordinate framework, SAMPLES make a crucial contribution in the ongoing modernization of brain biorepository practices.

By providing a spatially explicit, standardized atlas for brain biorepository resources, SAMPLES has the potential to improve harmonization across biorepositories, enhance biorepository workflows, streamline tissue requests, support precision neuropathology, simplify linking neuropathological findings with *in vivo* neuroimaging data, and facilitate collaboration between neuropathologists, imaging researchers, and molecular neuroscientists to unlock the research potential for research-relevant tissue samples in the study of neurodegenerative disease.

## Author Contributions

**JMW:** Conceptualization, Methodology, Software, Formal Analysis, Investigation, Validation, Visualization, Writing – Original Draft, Writing – Review and Editing.

**AS:** Writing – Review and Editing.

**YAS:** Formal Analysis, Software, Writing – Review and Editing.

**TL:** Writing – Review and Editing.

**ER:** Investigation, Supervision, Validation, Writing – Review and Editing.

**MB:** Investigation, Validation, Writing – Review and Editing.

**AK:** Investigation, Validation, Writing – Review and Editing.

**CMD:** Writing – Review and Editing.

**CSL:** Conceptualization, Resources, Investigation, Supervision, Validation, Writing – Review and Editing.

**CDK:** Conceptualization, Resources, Supervision, Funding Acquisition, Writing – Review and Editing.

**TJG:** Conceptualization, Supervision, Funding Acquisition, Writing – Review and Editing.

