## Supplementary Material for "Digital Atlases to Unlock the Potential of Brain Biorepository Tissues for Interdisciplinary Research"

Jason M. Webster[1*], Ali Shojaie[2], Yiqin Alicia Shen[1], Tung Le[2], Emily Ragaglia[3], Marika Bogdani[3], Amanda Kirkland[3], Christine Mac Donald[4], Caitlin S. Latimer[3], C. Dirk Keene[3], and Thomas J. Grabowski[1]

[1] Integrated Brain Imaging Center, Department of Radiology, University of Washington, Seattle, WA, USA

[2] Department of Biostatistics, University of Washington, Seattle, WA, USA

[3] Department of Laboratory Medicine and Pathology, Division of Neuropathology, University of Washington, Seattle, WA, USA

[4] Department of Neurological Surgery, University of Washington, Seattle, WA, USA

### Supplementary Figures

**A Routine Diagnostic Blocks**
A1: Right MFG
A2: Left MFG
A3: Bilateral prefrontal WM
A4: Right IPL at splenium
A5: Left IPL at splenium
A6: Right SMTG at ant comm.
A7: Left SMTG at ant comm.
A8: Right calcarine cortex
A9: Left calcarine cortex
A10: Bilat. anterior cingulate gyri
A11: Bilat. anterior hippocampus
A12: Bilat. mid hippocampus
A13: Bilateral amygdala and OB
A14: Right neostriatum at ant. comm. (with claustrum/capsules)
A15: Left neostriatum at ant. comm. (with claustrum/capsules)
A16: Right thalamus with STN
A17: Left thalamus with STN
A18: Midbrain w/ SN @red nuc
A19: Pons with locus ceruleus
A20: Medulla – mid level
A21: Bilat. cerebellum with cortex overlying dentate nucleus
A22: Spinal cord, pituitary gland, pineal gland
A23+: Gross lesions including infarcts, contusions, hemorrhages, mass lesions, etc.

**B Brainstem blocks**
B1: Rostral midbrain
B2: Midbrain
B3: Caudal midbrain
B4: Midbrain – Pons junction
B5-8: Pons
B9: Pontomedullary junction
B10: Rostral medulla
B11-12: Medulla
B13: Caudal medulla
B14: Rostral spinal cord

**C CTE blocks**
C1: Right SFG with SF sulcus
C2: Right SFG with SF sulcus
C3: Right anterior temporal lobe
C4: Left anterior temporal lobe
C5: Right orbital frontal cortex
C6: Left orbital frontal cortex
C7: Right hypothalamus
C8: Left hypothalamus
C9: Right MFG and sulci at SN
C10: Left MFG and sulci at SN

**D TBI blocks**
D1: Genu of corpus callosum (CC)
D2: Splenium of CC
D3: Body of CC at SN
D4: Body of CC with fornix
D5: Right middle cerebellar peduncle (MCP)
D6: Left middle cerebellar peduncle (MCP)

**E Neuroimaging blocks**
E1: Right medial prefrontal cortex
E2: Left medial prefrontal cortex
E3: Right hippocampus with PHS
E4: Left hippocampus with PHS
E5: Right precuneus
E6: Left precuneus
E7: Bilat post cingulate at LGN
E8: Right posterior angular gyrus
E9: Left posterior aangular gyrus
E10: Right fusiform/inferior TG
E11: Left fusiform/inferior TG
E12: Right posterior SMTG
E13: Left posterior SMTG
E14: Right lat parieto-occ ctx
E15: Left lat parieto-occipital ctx
E16: Right inferior frontal gyrus
E17 Left inferior frontal gyrus
E18: Right parietal WM at LGN
E19: Left parietal WM at LGN
E20: Right occipital WM
E21: Left occipital WM
E22: Right frontoinsula
E23: Left frontoinsula
E24: Supplementary motor cortex
E25: Right IFG – Broca
E26: Left IFG - Broca
E27: Right primary motor cortex
E28: Left primary motor cortex
E29+: Imaging-guided sampling

**Supplementary Figure 1**. The UW BRaIN Lab Modular Sampling Protocol for Precision Neuropathology. Sample are arranged in modules. The routine diagnostic module corresponds to the NIA-AA criteria for the assessment of Alzheimer's disease and related dementias. The brainstem module includes the entirety of the brainstem, though routine diagnostic samples take precedence (e.g. the B1 cassette is empty since this sample "is" A18). The chronic traumatic encephalopathy module is based on CTE consensus criteria. The traumatic brain injury module primarily contains white matter tracts often damaged through neurotrauma. The neuroimaging-guided module includes regions based on their role in functional networks, particularly those with disease-relevance.

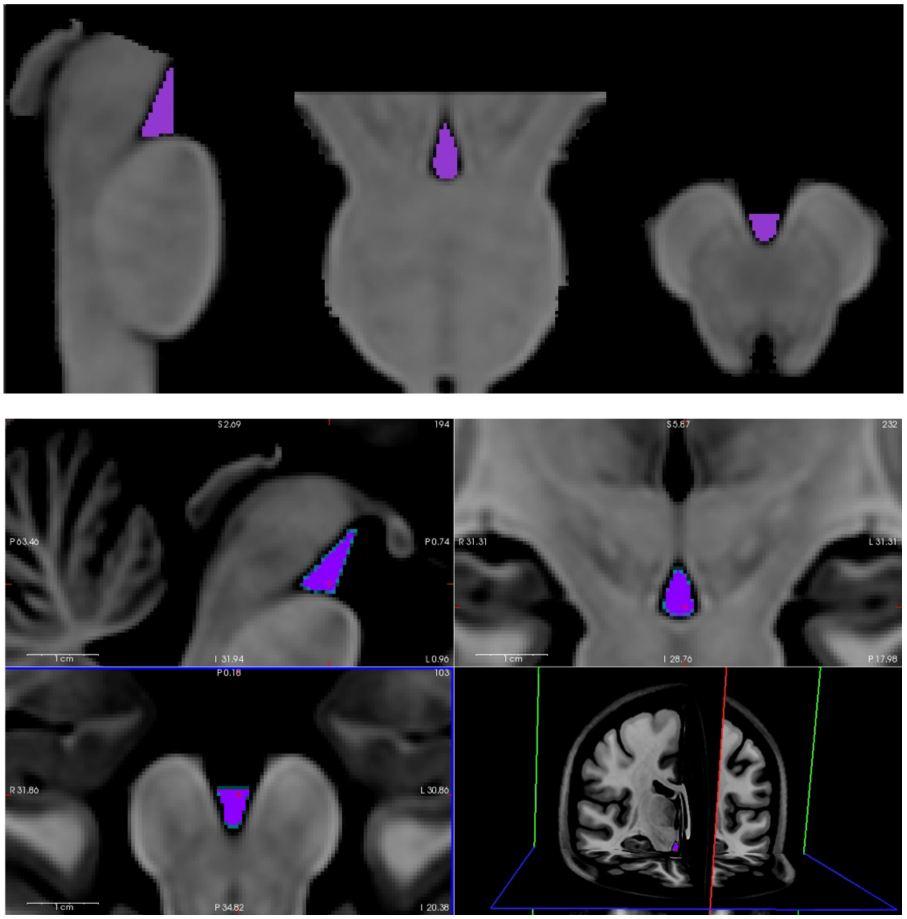


Supplementary Figure 2. Registration of the vertical brainstem volume. The UW BRaIN Lab slicing protocol takes 4 mm slices of the brainstem orthogonal to the major axis of the brainstem. The digital Brain Sampling Protocol atlas replicates this through axially slicing the vertical brainstem volume (top pannel) which was generated by applying an affine transform based on the principal component axis of the central sagittal slice of the cropped brainstem volume (not shown). The accuracy the approach was assessed by using the inverted affine to transform a manually created interpeduncular fossa label (purple) to the MNI2009b brain (bottom pannel).

### Supplementary Methods

The following sections provide additional detail on the procedures used to construct the digital Brain Sampling Protocol Atlas to precisely correspond to the UW BRaIN Lab's Brain Sampling Protocol.

#### Brain Extraction

Starting from the MNI2009b brain T1w volume, brain extraction was initialized with using FreeSurfer's *mri_synthstrip* (Hoopes, Mora, Dalca, Fischl, & Hoffmann, 2022) and then manually refined. In FreeSurfer's *freeview*, the highest value in this volume that appeared to be primarily CSF was determined at the interface with various tissue types at several locations. The minimum of these values was used passed to FSL's *fslmaths* to threshold the *brain extracted volume* to remove voxels below this value.

The *brain extracted volume* was then reloaded in *freeview* (color map= 'GE Color', min=0, max=95, opacity=.75). Next the MNI2009b brain T1w volume was loaded (min=0) and moved the layer below. Progressing superior to inferior, the polyline and flood fill tools were used to remove larger areas of non-brain voxels including skull, veins, and dura. Progressing through each orientation, any remaining non-brain tissue was removed with the freehand tool (brush value=0) and any omitted brain voxels were cloned (reference volume = MNI2009b brain). A final pass was made through each orientation, using the same approach for any refinements, to ensure all tissue boundaries were smooth and all brain tissues were complete.

#### Brain Segmentation

##### Brainstem Volume

An initial brainstem volume was created by identifying the coordinates of the bounding box of the brainstem using *freeview*, passing these values to FSL's *fslmaths* with the "-roi" flag to create a binary mask of the bounding box, and multiplying this with the brain extracted volume. This initial brainstem volume was then loaded into *freeview* and voxels containing the cerebrum and cerebellum were coarsely removed using the polyline and flood fill tools.

Following the UW BRaIN lab protocol for separating the posterior fossa from the cerebrum along the plane from the base of the mammillary bodies to the base of the thalamus, the polyline and fill tools were then used to remove voxels superior and anterior to the line between the mammillary bodies and base of the thalamus on each sagittal slice where the structures were clearly observed. Proceeding through the range of axial slices containing these structures, voxels to either side of the removed voxels were also removed. Then a final sagittal pass was made to remove all voxels superior and anterior the line connecting the removed voxels. A final axial pass was then made to remove any remaining cerebrum lateral or posterior to the brainstem.

Finally, a similar process was undertaken corresponding to the UW BRaIN Lab segmentation protocol for separating the brainstem from the cerebellum laterally along the curvature of the pons through the cerebellar peduncle and medially along the curvature the cerebellum through the cerebellar peduncles and medullary vermis.

##### Cerebellum Volume

An initial cerebellum volume was created by identifying the bounding box of the cerebellum in the brain extracted volume using *freeview*, passing these values to *fslmaths* with the "-roi" flag to create a binary mask, and multiplying with the brain extracted volume. Then *fslmaths* with the "-binv" and "-mul" flag was used to remove the brainstem from the initial cerebellum volume using an inverted binary mask. This volume was then loaded into *freeview* and the remaining fragments of cerebrum were manually removed, first with the polyline and flood fill tools, and then refined with the freehand and clone tools. The resulting cerebellum volume was then saved.

##### Cerebrum Volume

A cerebrum volume was then created by removing the brainstem and cerebellum voxels from the brain extracted volume. First, the cerebrum volume was initialized using *fslmaths* with the "-binv" and "-mul" flag was used to remove the brainstem from brain extracted volume. Then *fslmaths* with the "-binv" and "-mul" flag was used to remove the cerebellum from cerebrum volume.

#### Brain Slicing

##### Vertical Brainstem Volume

In the UW BRaIN Lab fixed-tissue BSP, the brainstem is sliced rostral to caudal along the long axis of the brainstem. In the MNI 2009b brain, the rostrocaudal axis of the brainstem is pitched forward relative to the axial dimension of the cerebrum, which is vertical in the voxel coordinate system. To enable uniform images corresponding to BSP's brainstem slices, the brainstem to the rostrocaudal axis, a custom **Python** script (ANTsPy + scikit-learn) was used to create a volume with the rostrocaudal axis of the brainstem positioned vertically. The rostral axis $\hat{v}_R$ of the brainstem was estimated from the *brainstem volume* as the first principal component of non-zero voxel coordinates from the central sagittal slice. The affine transformation for the minimal rotation angle $R$ about the volume centroid that aligns $\hat{v}_R$ to a vertical unit vector $\hat{z}$ of the voxel coordinate system ($R\hat{v}_R = \hat{z}$) was then applied to produce the *vertical brainstem volume* used in subsequent stages.

##### Initial brainstem slice

The initial brainstem slice will contain (A18: Midbrain with substantia nigra and red nucleus) is crucial for routine neuropathological diagnosis, an initial brainstem slice is taken parallel to the segmentation cut to ensure thickness and inclusion of the relevant neuroanatomical structures.

In *the vertical brainstem volume*, the anatomically determined segmentation cut is a slightly anteriorly tilted plane. This plane was computationally determined by identifying the most superior horizontal slice with nonzero voxels, setting the average location of those voxels as a pivot point, finding the unit vector orthogonal to the plane, and iteratively tipping the plane about the pivot until it contacted the segmentation cut. The *initial brainstem slice image* in png format was created from the voxels intersecting rotated plane. The coordinate frame of this plane centered on the pivot point was used to identify the cutting plane 4 mm below for the initial brainstem slice. Voxels corresponding to sample A18 were identified through positioning a rectangular mask matched to the sampling tool dimensions on the segmentation plane and 'cutting' through the slice, then copying the label these voxels into the *brainstem sample volume*. The initial brainstem slice was then removed from the *vertical brainstem volume*.
